# Temporal Trends: Phase-shifted time-series analysis reveals highly correlated reproductive behaviors in the black soldier fly, *Hermetia illucens* (Diptera: Stratiomyidae)

**DOI:** 10.1101/2025.08.26.672371

**Authors:** Noah B Lemke, Chujun Li, Jeroen De Smet

## Abstract

For those researching reproduction of the black soldier fly (BSF), finding meaningful correlations between mating and downstream reproductive fitness metrics, owing to complex reproductive behavior and physiology, has been challenging. On the one hand, mating is not necessarily deterministic of fertile eggs being produced, since females might lay unfertile eggs both before and after mating. But conversely, the process is not ‘random’ either, since fertility is predicated on the exchange of gametes during mating. To investigate the relationship between mating and other reproductive metrics, a cross-correlation analysis between lagged time series was conducted on data previously collected during a semi-outdoor greenhouse experiment. This process compared the average percent correlation between un-lagged mating with oviposition in traps, oviposition outside traps, total weight of eggs deposited, and hatch rate, which were lagged 0 to 6 days. This revealed that average mating was 97.54% correlated with oviposition in traps lagged by 2 days. For oviposition outside of traps, total egg weights, and hatch rate, these were also ∼71.63% correlated, 60.29% correlated, and 75.16% correlated with mating respectively, which were optimized when the time series were lagged n = 3 days. These trends likely have a biological basis since flies decrease their reproductive activity as they expend energy and senesce, aligning with temporal patterns consistently observed during BSF rearing.

## INTRODUCTION

Insect reproductive behavior is complex (Lemke et al. 2023, Meneguz et al. 2023, Tomberlin et al. 2025) owing not only to the wide diversity in mating systems and reproductive modes across species (Kokko et al. 2014, Normark 2014), but also the plethora of alternative mating tactics, dynamic fitness optima, behavioral phenotypes, and personas (Shuker and Simmons 2014). As an emergent phenomenon (Macklem 2008), reproductive behavior is influenced by interactions occurring at lower levels of biological organization. These range from perceptions of group-social context, hormonal state, physiology, nutritional legacy, cellular processes, genetics, eco-evolutionary history (Alcock and Thornhill 2014) and much more. On top of this, studies of mating behavior must also consider non-biological factors that can influence behavioral outcomes and their interpretation, such as the testing environment, experimental artifacts, and observer biases (Tomberlin et al. 2025). As such, a broad intuition of insect biology as well as sophisticated analytical techniques are needed to understand behavioral complexity (Smetana et al. 2025).

The black soldier fly (BSF), *Hermetia illucens* (L.) (Diptera: Stratiomyidae) is a large lekking fly (Tomberlin and Sheppard 2001, Lemke, Smith, et al. 2025) of neotropical origin (Kaya et al. 2021) with complex mating behavior. BSF has been recently domesticated (Caruso et al. 2013, Generalovic et al. 2023) and is now mass-reared throughout the globe for various applications, such as waste management and subsequent conversion of these materials into insect biomass that can serve as alternative feed ingredients for pets and livestock. For this reason, the operation costs of mass-rearing facilities still need to be driven down to make facilities more economic since the goal of a mass-culture program is to produce the maximum number of fertile females in as short of time and inexpensively as possible with minimal labor and space (Boller 1972).

Due to protandry, adult male BSF emerge from their pupae prior to females (Tomberlin et al. 2002, Generalovic et al. 2025). Mating studies typically control for age-sex distributions of emerging adults by taking a portion of reared flies and sorting them into test populations (e.g., 100 males and 100 females). This approach can be contrasted with the industrial process, in which one (or several) batch of pupae are placed directly inside a cage, or in an area adjacent to the cage and allowed to emerge (Coudron et al. 2025). If multiple batches of pupae are added to the cage continuously, then this can lead to the breeding cage containing flies of many mixed ages and reproductive statuses (Dickerson et al. 2024, Lemke, Li, et al. 2025) especially since post-reproductive adult flies might live several weeks or more (Harjoko et al. 2023) when supplemented with water and food (Thinn and Kainoh 2022, Klüber et al. 2023, Barrett et al. 2025).

Irrespective of the exact method used to stock cages, what follows is that high densities of individuals engage in ‘lek-like’ mating (Lemke, Smith, et al. 2025) which typically peaks 1-3 days after being introduced to the breeding environment (Dickerson et al. 2024, Lemke, Li, et al. 2025). Males are drawn towards UV-AB and blue-green light (Zhang et al. 2010, Oonincx et al. 2016), and females temporarily join these aerial swarms to mate (Lemke et al. 2024). After an aerial courting ritual, copulation commences when male and female lock genitalia and remain nearly motionless on the floor or walls of the cage for ∼35 minutes (Manas et al. 2025). Near the end of this time, sperm is transferred via the male’s aedeagus into the female (Manas et al. 2025), the two separate, and groom themselves (Laudani et al. 2024).

Following this, there is an intermittent period of several days (Tomberlin and Sheppard 2002) where sperm travels through the female’s genital tract (Malawey et al. 2019) and is stored in their spermathecae until the moment of oviposition (Manas et al. 2024), whereupon the eggs are fertilized. To induce egg-laying, an attractant is then introduced to entice females to lay their eggs in a trap. Oviposition activity typically then plateaus between days 5-6 days after being introduced into the breeding environment (Dickerson et al. 2024). To lay their eggs, females normally seek out an area of dry cracks adjacent to a larval rearing substrate (but obviously such a substrate may not actually be present if the attractant is a synthetic blend). Traps are then replaced, and any deposited eggs are harvested by technicians. 2-5 days later, the neonates hatch from their eggs and begin wandering towards food (Giannetti et al. 2022).

One major area complicating analysis is the temporal structure of the data. Mating and oviposition activity peak several days apart (Tomberlin and Sheppard 2002, Dickerson et al. 2024, Meyermans et al. 2025). Moreover, outside of these peaks there is very little activity for cohorts that have homogenous underlying age structure, leading longitudinal count data to be highly zero-inflated (Smetana et al. 2025). To capture information about both mating and oviposition, observational experiments need to be run for several days (Chiabotto et al. 2024, Dickerson et al. 2024, Lemke et al. 2024), which presents a logistical hurdle because, unless remote videography is used, human observations cannot accurately be made across this whole time period. A trade-off then exists between capturing discrete observations across the entire cycle (e.g. during daylight hours for 1 week) or focusing continuously on a narrower time period (e.g., the time-windows for peak mating or oviposition). Furthermore, the temporal delay between copulation (i.e., mating), oviposition, and eggs hatching has historically made predicting the values of one from the others difficult, leading some reports to say that mating is only loosely related to the latter (Dickerson et al. 2024).

Hence, the objective of this study is to demonstrate the relationship between mating and oviposition, and whether mating could be used as an upstream predictor for fertile egg production. This study examined whether the correlation between BSF mating and other reproductive metrics could be improved through phase shifting (i.e., lagging) their time series. While the procedures for time series analyses are formalized and widely used in field studies of insect populations (Hoshi et al. 2014, Poh et al. 2019, Chaves and Friberg 2021), as well as other fields like sociology and economics (Shumway and Stoffer 2017a), such an analysis is new when it comes to interpreting black soldier fly reproductive behavior. We hypothesized that by adjusting the lag between time series, that mating could be a useful predictor of downstream fitness metrics such as oviposition, total egg weight, and hatch rate.

## METHODS

### Data Acquisition

Data for this analysis was originally collected as part of a semi-outdoor greenhouse study conducted at the F.L.I.E.S. Facility at Texas A & M University throughout 2022 (Lemke, Li, et al. 2025). The original experiment tested the effect of oviposition attractant provisioning on fitness as well as higher-order interactions between BSF cohort age, attractant provisioning, and fitness. It did not, however, consider the possible temporal relationships between fitness variables (i.e., mating, oviposition, total egg weight, and hatch rate), which is done here via a time-series correlation analysis.

In the original experiment, data were collected during four trials that took place from (a) March 07-13, (b) April 25-May 01, (c) October 06-12, and (d) December 10-16, each lasting seven days. During these observation periods, count data for mating and oviposition were recorded hourly from sunrise to sunset, while eggs traps were collected daily from cages at noon and the total weight of collected eggs measured on an electronic balance scale. After a five-day incubation period, hatch rate of the eggs for each collection was also assessed. A full description of the experimental design can be found in Lemke et al., 2025; and similar procedures have been used in other studies (Dickerson et al. 2024, Laursen et al. 2024, Lemke et al. 2024). During each trial, four experimental treatments (see below) were tested on the production of a 0.93-m3 breeding cage, supplied with 100 male and 100 female BSF. Each of these four treatments was replicated three times, for a total of twelve experimental units, yielding 336 cage-days of observations (4 treatments × 3 replicates × 2 blocks × 2 trials × 7 days).

### Data Structure & Cleaning

The levels of the original experiment’s treatment categories followed a 2 × 2 factorial design, with attractant provisioning having four total combinations of levels (i.e., (a) whether the attractant provided initially or (b) delayed until 1-days post peak mating; and (c) whether the attractant contained substrate or (d) was an empty control). Additionally, age group also had an additional two blocks (two trials each, of which all flies were either “mixed” or “same” age). For the age groups, “young” flies were 1-4-day-old prior (viz., 2-4-day-old males, and 1-3-day-old females) to their introduction into mating cages, whereas the “mixed” cohort could have ages between 1-16-days.

However, because industrial producers of BSF do not delay the age of their flies prior to release, for this analysis, we only considered data for the “young” treatment in which this artifact was absent. Likewise, providing attractant boxes lacking attractant is also not representative of industrial rearing conditions and was included in the original experiment as a negative control, but was not taken into consideration for this analysis. Moreover, recent evidence has shown that virgin females have no inherent preference towards a group of volatile organic compounds (VOCs) that later caused attraction or avoidance behaviors in mated females (Klüber et al. 2024). And so, in addition to “mixed” age flies, replicates for which there was never an attractant provided (labeled as “IC” and “DC” respectively for Initial and <u>D</u>elayed <u>C</u>ontrols) were redacted from the data set. This yielded 168 rows of data (2 trials × 12 cages × 7 days) (i.e., half of the original 336 cage-days). The reduced data frame (containing only data from “young” flies that were provided attractant initially = “IO” or delayed “DO”) was then reformatted using the PivotTable function in MS Excel for Mac Version 16.97 (25051114), to make calculating the averages much easier.

### Time Series Analysis

To examine temporal patterns, the data were plotted using MSExcel inbuilt plotting function. By averaging values by day (of observations taken hourly from sunrise to sunset) autoregression (as used by Lemke et al. 2024) was not necessary for this analysis because there were no gaps in the data that needed to be filled. Visually inspecting the plots revealed that phase shifting the data might lead to stronger cross-correlations, since mating and oviposition follow a similar temporal pattern, albeit with oviposition lagging by two days.

### Percent Correlation Coefficient Matrix

To make cross-correlations, a Pearson correlation matrix was constructed which included all pairwise combinations for mating and oviposition across treatments. For example, the mating-oviposition (in traps) matrix considered the mean average of IO-mating, as correlated to the mean average of DO-mating, IO-oviposition, and DO-oviposition. For the resulting matrix, only the lower triangle of correlation coefficients (i.e., cells underneath the identity diagonal) were considered since the matrix was symmetrical. Values were then squared (r-squared), which is also considered a measure of effect size (Rights and Sterba 2020), and multiplied by 100 to yield the Pearson Percent Correlation Coefficient (PCC). From this, the average Pearson PCC of the matrix was calculated (i.e., by summing all values and dividing by the number of values within the lower triangle), as well as the standard deviation and standard error (of n = 6 values under the diagonal). For this step, Pearson’s correlation was chosen to investigate when a linear increase in mating corresponded to a linear increase in oviposition, or vice versa. However, using Spearman’s rank correlation would allow a looser interpretation in which an increase in mating corresponded to *any* increase in oviposition, not necessarily a linear one (and was used in the Clustered Heat Map analysis below).

### Phase Shifted PCC

Next, a series of Pearson PCC matrices were calculated to determine what amount of phase shifting maximized average Pearson PCC. This was done for each pair of reproductive metrics (mating with oviposition in traps, mating with oviposition in cages, mating with total egg weight, and mating with hatch rate). Data were phase shifted n = 0, 1, 2, 3, 4, 5, and 6 days, yielding *y_t_, y_t-1_, y_t-2_, … y_t-6_*. To phase shift the data, the first n-days of the data were deleted and the remainder shifted forward, creating an empty tail that was filled with 0-values to preserve the length of the time series. Mating was always kept unmodified, primarily because this behavior generally occurs first in the reproductive sequence (barring instances where mating is delayed for several days), which means a oviposition phase shift as a function of mating is the more biologically relevant parameter combination to explore rather than vice versa. The average Pearson PCC across this continuum of phase shifting was then plotted in MSExcel and examined visually to determine candidates for optimization by finding local maxima/minima. Lastly, to determine the potential for an underlying autoregressive structure (i.e., temporal non-independence) to bias the average Pearson PCC scores (Shumway and Stoffer 2017a), the partial autocorrelation function (PACF) of mating versus lagged mating was computed via the pacf() function in R. Doing so revealed that none of these lags were statistically significant, i.e., PACF scores falling above or below 95% confidence intervals, suggesting temporal independence (Supplementary Figure 1).

### Clustered Heat Map

Finally, a clustered heat map was generated for all metrics, with treatment data pooled together. The input data frame for the heatmap was constructed using reproductive metrics lagged however much would maximize average Pearson PCC, as well as their neighboring lags of ± 1 day (i.e., if an optimum was found for oviposition with a phase shift of n = 2, then this along with n = 1 and n = 3 were considered). The correlation matrix was calculated using the corr() function in RStudio (Wei and Simko 2024), using Spearman’s rank coefficient (since the functions above were not strictly monotonic). These values were then plotted as a clustered heatmap using the corrplot() function in RStudio, using Euclidean distance to construct the distance matrix, and then ordered using a single-linkage hierarchical clustering algorithm (which builds trees via a bottom-up agglomerative approach).

## RESULTS

### Visual Analysis

As a first step, data for mating × day, oviposition × day, total egg weight × day, and hatch rate × day was visualized by plotting in RStudio (Figure 1-A, Figure 1-B). Doing this revealed average mating peaked on day 1 of the experiment, (i.e., after flies were introduced into the breeding environment), whereas average oviposition peaked on day 4. The trend for each was to generally decrease following this peak, however mating was not strictly monotonic, in that there was a second small peak between days 3-4. Cumulative total egg weight and hatch rate both plateaued by day 5, with values generally decreasing each day after reaching a peak on day 3. Qualitatively, it then appeared that mating would have the strongest correlation to oviposition when the latter was lagged n = 2 days. However, because of the plateau, total egg weight and hatch rate might be more strongly correlated with mating when phase shifted greater than 1 day.

**Figure 1.**
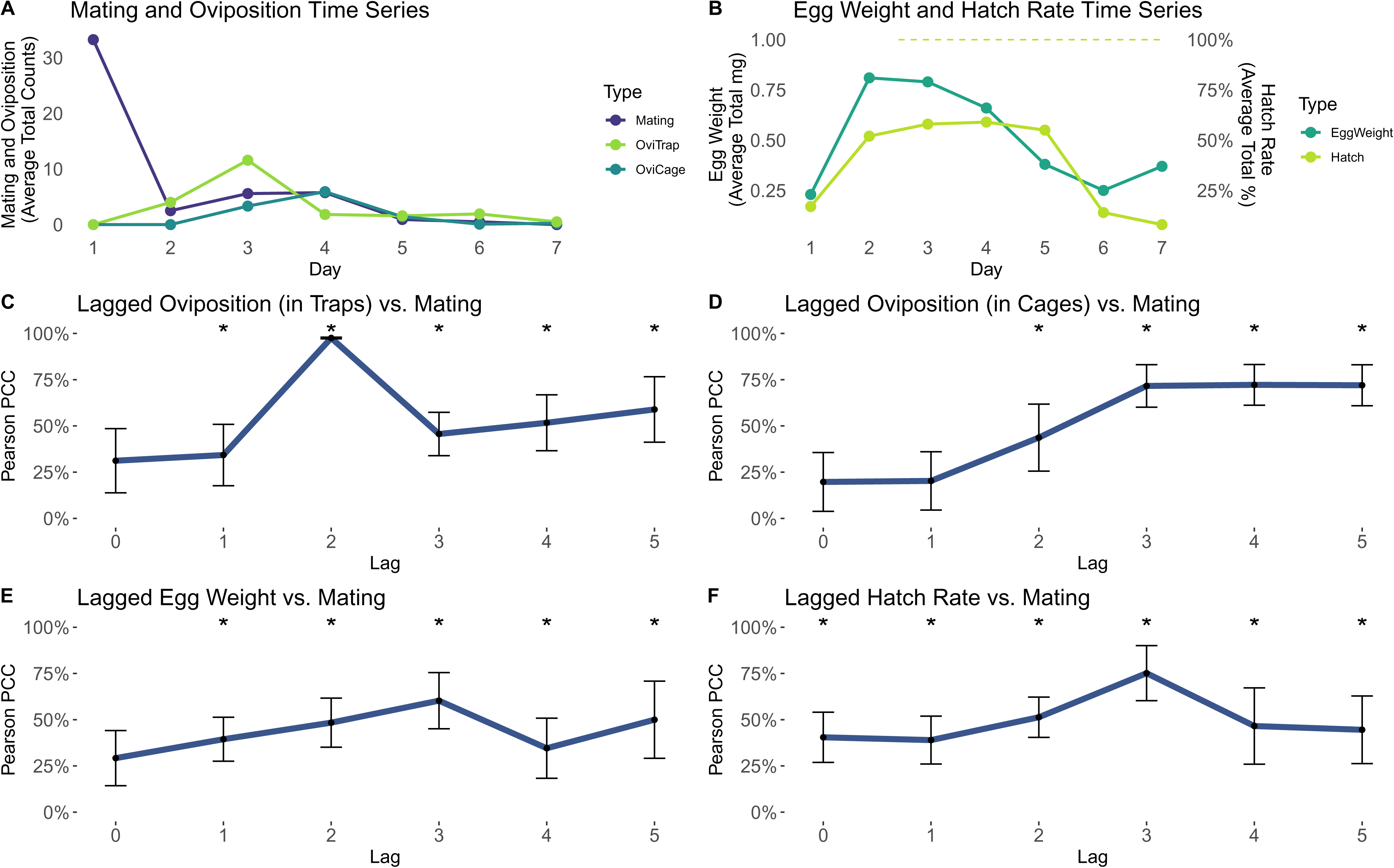
Time-series data for BSF fitness metrics and correlations between lagged BSF fitness metrics and un-lagged BSF mating counts. (A) Time series of mating counts and oviposition counts. Data was collected from twelve, 0.93-m3 cages each supplied with 100 male and 100 female adult black soldier flies aged <4 days. Hourly observations were made from sunrise to sunset for seven days, and for two trials. Counts were summed per day and then averaged across cage/trial to yield average total counts with respect to each day. (B) Time series data for total egg weight and hatch percentage. Eggs were harvested from traps once per day, and the total weight was averaged across cages. For each collection, hatch rate was also measured, and these rates were likewise averaged across cages to yield the average percent hatch rate. (C) Average Pearson Percent Correlation Coefficients (Pearson PCCs) for oviposition in traps lagged 0-5 days against un-lagged mating counts. (D) Average Pearson PCC for oviposition in cage material lagged 0-5 days against un-lagged mating counts. (E) Average Pearson PCC for total egg weights lagged 0-5 days against un-lagged mating counts. (F) Average Pearson PCC for hatch rates lagged 0-5 days against un-lagged mating counts. For plots C-F, error bars indicate ± SD. Average Pearson PCC for lags of 6 days have been excluded from these plots as outliers. FIGURE 1 ALT TEXT: Multi-panel plot of time-series data for BSF reproductive behaviors, as well as the average Pearson Percent Correlation Coefficient (PCC) between mating and lagged reproductive metrics. Panel A depicts average mating and oviposition over 7 days. Panel B depicts average total-egg weight and hatch rate over 7 days. Panels C-E depict average Pearson PCC for mating vs. lagged oviposition (in traps), mating vs. lagged oviposition (in cages), mating vs. lagged total egg weight, and mating vs. lagged hatch rate; respectively, for lags of n = 0, 1, 2, 3, 4, and 5 days.

### Cross Correlation of Time Series Data

The initial analysis considering Mating × Oviposition (in traps) for each treatment was optimized when oviposition was lagged by 2 days, with individual values within the matrix ranging from 97.56 - 98.66 average Pearson Correlation Coefficient (PCC) (Table S2), indicating that PCC could be optimized by lagging data. Hence, for each reproductive metric, the average Pearson PCC was calculated between mating, and that second metric phase shifted by n = 0, 1, 2, 3, 4, 5, and 6 days. These were then plotted to visually inspect where (local) optima lie. For the comparison between mating and oviposition (in traps), average PCC was maximized with a phase shift of n = 2 days. Doing this increased the average Pearson PCC from 31.19% to 97.54%, a 3.12-fold difference (Figure 1-C). For the comparison between mating and oviposition (in cage material) (Figure 1-D), average Pearson PCC was maximized with a 3-, 4-, or 5-day time lag (∼72%) and minimized with a 0-day time lag (19.71%). For the comparison between mating and total egg weight (Figure 1-E), a local maximum was found with a 3-day lag (60.29%) up from 29.22% with no lag, representing a 2.06-fold increase. Lastly, the comparison between mating and hatch rate (Figure 1-F) also had a maximized average Pearson PCC with a 3-day lag (75.16%) up from 40.48%, representing a 1.85-fold increase. Interestingly though, often the highest value, regardless of the metric being examined was a 6-day time lag, though this is suspected to be a statistical artifact (see Discussion below). Interestingly, ∼98-99% correlations were found between mating and metrics phase-shifted n = 6 days, but these may be a statistical artifact (i.e., a spurious correlation) from comparing time series whose tails were mostly filled with zeroes, and so these were discounted as outliers.

### Clustered Heat Map

A clustered heat map was constructed using single-pass hierarchical clustering algorithm to reveal grouped relationships between mating and downstream reproductive metrics based on the Euclidean distance of their Spearman rank correlations (Figure 2). Input variables that went into this heat map included (a) unmodified fitness metrics, (b) metrics phase-shifted an amount equal to the optima found in the previous step, and (c) phase shifts of ±1 days from the optima. Within the clustered heat map, mating was most similar to oviposition in traps phase shifted 2 or 3 days as well as total egg weight and hatch rate each shifted 3 days. Mating was moderately similar to oviposition in traps shifted by 1 day as well as total egg weight and hatch rate shifted by 4 days. Mating was most dissimilar to a cluster containing oviposition in cages (phase shifted 0, 2, 3, and 4 days), as well as a cluster containing all the unmodified metrics.

**Figure 2.**
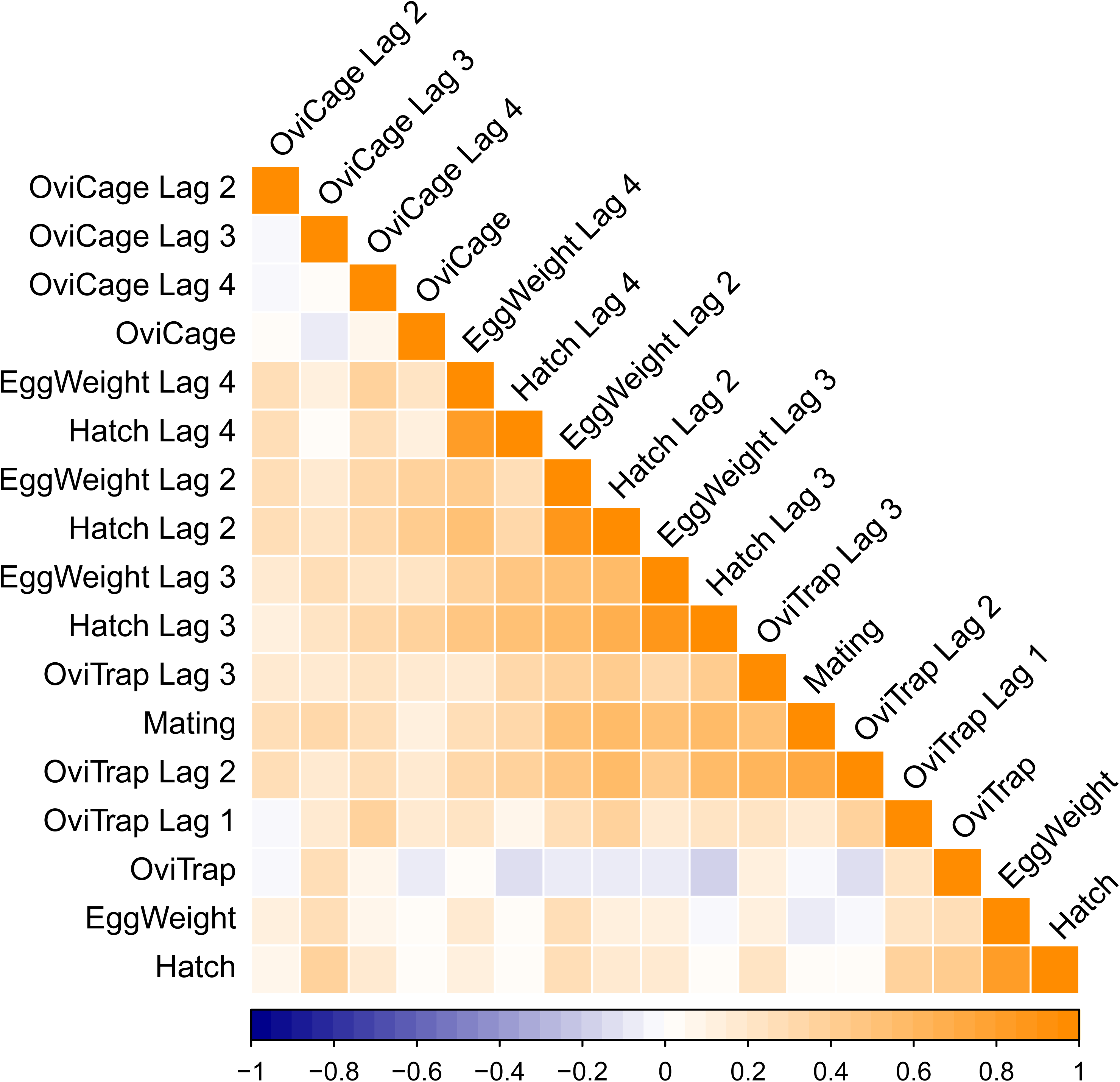
Clustered Heat Map of Time Series Data. Correlation matrix was calculated using Spearman’s rank coefficient. Distance matrix was calculated using Euclidean distance. Ordering was performed using single-pass hierarchical clustering algorithm. Abbreviated data labels indicate the following: “Mating” = Mating. “OviTrap” = Oviposition in traps. “OviCage” = Oviposition in Cages. “EggWeight” = Total Egg Weight. “Hatch” = Hatch rate. FIGURE 2 ALT TEXT: Clustered heat map indicating that the Spearman’s Rank correlation between mating and the other reproductive metrics is highest when those other metrics are phase shifted 2 days for oviposition in traps, between 2 and 4 days for egg weight, and 3 days for hatch rate. Mating is most distant to non-shifted metrics as well as oviposition in cage material regardless of amount the latter is phase shifted.

## DISCUSSION

The objective of this study was to examine whether correlations between time series data of reproductive metrics recorded during a BSF breeding study could be improved by comparing time series that were phase shifted/lagged. Since there is an obvious temporal component to mating behaviors, this study improves upon that of Lemke, Li, et al. 2025 which did not consider temporal interactions as part of the original study, focusing instead on statistical differences between treatment effects. Indeed, the results of this analysis show that lagging time series could increase the Pearson percent correlation coefficient (PCC), i.e., effect size, between mating and other downstream reproductive metrics substantially. By performing this transformation, mating was 97.56 to 98.66% correlated with oviposition in traps if lagged by 2 days. Likewise, for oviposition outside of traps, total egg weights, and hatch rate, these were ∼71.63% correlated, 60.29% correlated, and 75.16% correlated with mating respectively if their time series were lagged n = 3 days. A clustered heat map confirmed this relationship, showing mating was most closely related (i.e., had the closest distance) with a cluster of metrics that included oviposition, total egg weight, and hatch rate, that were each phase shifted by these optima. In addition, the clustered heat map showed that mating was also most different from these same fitness metrics if they were unmodified and was also unrelated to oviposition in cage material. Thus, this experiment highlights the importance for BSF producers to considering correlations between reproductive metrics measured at different time points in production, as well as understanding whether these correlations are rooted in biology.

At first glance, the link between mating and these lags generally corresponded biologically to temporal patterns within the data. The initial visual inspection of the reproductive metrics (plotted by day) revealed that mating peaked on day 1 of the experiment (Figure 1). The trend for the mating time series was then generally decreased but had a small ‘hump’ on days 3-4 (Figure 1). Because the underlying age structure of this data was relatively homogenous (1-4 d-olds), this suggested that individuals can engage in multiple matings, though it is certainly possible that small, but uneven differences in mating proclivity could also produce this pattern. That said, both male and female BSF have been verified by many recent experiments (Hoffmann et al. 2021, Jones and Tomberlin 2021, Chiabotto et al. 2024, Muraro et al. 2024) to mate multiple times. Similarly, oviposition in traps peaked on day 3 and then decreased monotonically (Figure 1), meaning the time difference between peak mating and peak oviposition (occurring on days 1 and 3, respectively) was n = 2 days, which was equivalent to the optimized time lag.

Interestingly, the peak for total egg weight occurred prior to the peak for oviposition counts in traps (Figure 1-A), indicating that perhaps larger egg clutches are laid by females earlier in the cycle, and smaller ones later in the cycle (or perhaps an incomplete initial oviposition). A prior experiment reports multiple oviposition events, but that subsequent clutches were smaller and infertile (Nakamura et al. 2016). Alternatively, it could also mean that later in the cycle there are instances where females extend their ovipositors but do not lay eggs (reflecting an inaccuracy in the original data set). Lastly, hatch rate peaked on day 3 (i.e., eggs laid on day 3 had the highest hatch rate after their 5-day incubation period). The hatch rate then decreased monotonically, suggesting that the process of sperm storage within the female takes ∼2 days (since before day 3 the hatch of any eggs collected is relatively low), aligning with a recent report (Manas et al. 2024). Alternatively, because the data were gathered in an semi-outdoor environment, it could also be that this peak in fertility corresponded to days with more optimal environmental conditions, such as higher humidity that might reduce desiccation, or ambient temperatures falling within thermal tolerances (Chia et al. 2018, Addeo et al. 2022).

Another interesting pattern in the data is that the average Pearson PCC between mating and oviposition (inside of traps) peaked earlier than did the average Pearson PCC between mating and oviposition (inside of cages), which seem to correspond to a 1-day earlier peak in oviposition in traps versus cages (Figure 1-B). Since the data used here include only 1-4-day-olds, the uptick in oviposition outside of cages can’t easily be explained by delayed mating, as was the case for prior experiments which did this to create aged test-populations (Dickerson et al. 2024, Lemke, Li, et al. 2025). Instead, the pattern might indicate that the flies become less discriminating as they age. There could also be an environmental effect, whereby the temperature happened to increase on those days (for both trials), forcing the flies away from ovipositing in the trap, which absorbed/reflected a lot of heat (Lemke, Li, et al. 2025). Another explanation could also be a snowball-effect whereby day 4-5 of the experiment, some flies had haphazardly laid eggs in the cage material early on, and then the volatiles from those eggs (Klüber et al. 2023) attracted additional flies to lay outside of the trap.

The temporal trends reported here generally align with what has been observed in previous BSF breeding experiments (Tomberlin and Sheppard 2002, Zhang et al. 2010). For example, early experiments on BSF showed that mating peaked on days 1-2, with oviposition peaking several days later (Tomberlin and Sheppard 2002, Park et al. 2010). This has been a consistent trend in mating experiments to this date (Jones and Tomberlin 2021, Dickerson et al. 2024, Lemke et al. 2024, Meyermans et al. 2025). In addition, more recent studies have demonstrated multiple mating (Hoffmann et al. 2021, Chiabotto et al. 2024, Muraro et al. 2024), multiple oviposition (Muraro et al. 2024), and physiological processes occurring during these ‘lulls’ in activity (Manas et al. 2024), as well as the potential for a neurophysiological switch during this time for females to become receptive to volatiles (Klüber et al. 2024). As such, it is reassuring to know that these patterns continue to hold true after two decades of BSF rearing, both throughout the globe, as well as in indoor and semi-outdoor rearing conditions.

However, because the original trend for mating in this experiment was to generally decrease after day 1, the average Pearson PCC between mating and any other reproductive metric appears to have been optimized when the phase shift applied also transformed the time series to become a monotonic decreasing function. Of course, this is reflective of the fact that the original experiment only used a single batch of flies per cage, and it is expected that this trend will not hold true once new flies are added to cages through continuous rearing (see below for suggested analysis to perform in this case). For the batch-reared data analyzed here, although mating was not strictly monotonic, in general its trend was to decrease.

Lagging the data n = 1 days effectively removed the peak on day 1 and made the new mating peak occur between days 3-5 (shifted to days 2-4). Hence, autocorrelation was highest on days 3-5, because lagging the function by this amount effectively transformed it into a strictly decreasing (i.e., monotonic) function. Likewise, the average Pearson PCC between mating (unadjusted) and total egg weight was highest when egg weight was lagged by 6 days (creating a strictly decreasing function to qualitatively match mating). Still, these trends have a biological basis that resonates with long held understanding of temporal patterns in BSF reproduction, in which reproductive activity decreases with time as flies exhaust their energy stores, and senesce (Harjoko et al. 2023, Dickerson et al. 2024).

Of course, correlation does not imply causation, but to some extent we can infer directionality since mating generally occurs prior to oviposition, oviposition before any eggs are collected, and total egg weight necessarily occurring before any are hatched. In addition, all these quantities intuitively must co-vary with one another, given that they are predicated on optimized abiotic conditions (e.g., light, temperature, humidity).

### Limitations and how to adapt this study for Industry

This analysis is limited largely because it uses a dataset collected from two disjunct rearing cycles only lasting 7 days each. For industrial programs that might need to produce high volumes of eggs/neonates, this analysis could be applied but would need to be modified in one of the following ways:

First, for those practicing batch rearing (Coudron et al. 2025, Lemke, Li, et al. 2025), mating might instead peak on day 2 or 3 of a breeding cycle rather than day 1 which would make sense due to differences in the emergence time between males and females (i.e., protandry) or their initial period of inactivity. In this case, since monitored mating activity would increase suddenly and then decrease, the mating time series would no longer be monotonic, and might be qualitatively similar to some of the other reproductive metrics. Similarly, the time series might be non-monotonic if BSF experience several activity peaks as a result of multiple mating. In either case, a Spearman’s Rank correlation could be calculated instead of Pearson’s to loosen some of the assumptions and examine increases in non-monotonic functions, which is what was done to construct the clustered heat map presented here (Figure 2).

If instead, a producer operates their breeding cages continuously (Dickerson et al. 2024, Coudron et al. 2025, Lemke, Li, et al. 2025), then it might be better to calculate a rolling window (i.e., a fixed time frame of e.g. 7 days, calculated each day) from which long term temporal patterns could be ascertained (such as, whether mating on certain days is correlated with oviposition on others, or whether there are weekly and seasonal trends across operations). Such patterns could be also modeled arithmetically through an ARIMA model (Shumway and Stoffer 2017b), which are traditionally used in stock price and commodities forecasting. Application of this technique will be complex however if there is a non-stationary trend or autocorrelation between rearing cycles (i.e., decreases over time due to genetic deterioration, or increases corresponding with sudden increases in population density).

The extent to which these are practical to implement for BSF breeding will of course depend on the elegance of behavioral assays and the information gleaned from them (Tomberlin et al. 2025). Standard metrics like egg weight and hatch rate, which are simple enough to measure gravi-, or volu-, metrically, can of course be formatted as Time Series data as has been done here. In addition, there are also many remote detection methods proposed for use in insect agriculture (Manoukis and Collier 2019, Nawoya et al. 2024), and some have started to be realized. These include FlyCount (James et al. 2024), a program that monitors population density of adults passing through a narrow opening prior to entering the breeding cage, as well as HistoNet (Sharma et al. 2020) to automatically assign size-classes to BSF. Counting the number of flies in a cage, recording their sizes, and monitoring fertile egg production in relation to these factors longitudinally (Dickerson et al. 2024) will be one step, but an important one, towards understanding the system and increasing its predictability in a complex system.

## ACKNOWLEDGEMENTS

We would like to thank our anonymous reviewers, as well as the group of researchers who helped collect the original data set: Amy J. Dickerson, D. A. Salazar, J. Emmanuel Mendoza, Lisa N. Rollinson, Erin Harris, Sienna McPeek, Rachel MacNeal, and Drs. Chelsea Miranda, and Casey Flint.

This material is based upon work supported by the National Science Foundation Graduate Research Fellowship under Grant No. 1746932. Any opinion, findings, and conclusions or recommendations expressed in this material are those of the authors and do not necessarily reflect the views of the National Science Foundation. This research was also partially supported by funds from the US State Department and the Belgium-Luxembourg Fulbright Commission under the 2026 US-Belgium Fulbright Scholar Program. The content of this report is explicitly that of the authors and does not reflect the views of the Fulbright Program, nor the governments of the United States, Belgium, or Luxembourg. This research was also supported by funds from the F.L.I.E.S. facility at Texas A&M and with material purchased from EVO Conversion Systems, LLC (Bryan, Texas, USA).

## REFERENCES

Addeo NF, Li C, Rusch TW, et al. 2022. Impact of age, size, and sex on adult black soldier fly Hermetia illucens L. (Diptera: Stratiomyidae) thermal preference. Journal of Insects as Food and Feed. 8(2):129–139. 10.3920/JIFF2021.0076

Alcock J, Thornhill R. 2014. The evolution of insect mating systems. In: Shuker DM, Simmons LW, editors. Evolution of Insect Mating Systems. Illustrated. United Kingdom: Oxford University Press. p. 78–91.

Barrett M, Patel N, McCarry B, et al. 2025. Dietary preferences and impacts of feeding on behavior, longevity, and reproduction in adult black soldier flies (Diptera: Stratiomyidae; *Hermetia illucens*). Journal of Insects as Food and Feed. Online:1–12. 10.1163/23524588-bja10295

Boller E. 1972. Behavioral aspects of mass-rearing of insects. Entomophaga. 17(1):9–25. 10.1007/BF02371070

Caruso D, Devic E, Subamia IW, et al. 2013. Technical handbook of domestication and production of Diptera black soldier fly (BSF), *Hermetia illucens*, Stratiomyidae.

Chaves LF, Friberg MD. 2021. *Aedes albopictus* and *Aedes flavopictus* (Diptera: Culicidae) pre-imaginal abundance patterns are associated with different environmental factors along an altitudinal gradient. Current Research in Insect Science. 1:100001. 10.1016/j.cris.2020.100001

Chia SY, Tanga CM, Khamis FM, et al. 2018. Threshold temperatures and thermal requirements of black soldier fly *Hermetia illucens*: Implications for mass production. PLOS ONE. 13(11):1–26.

Chiabotto C, Grosso F, Doretto A, et al. 2024. Observation of mating behavior using marked flies of black soldier fly (*Hermetia illucens*) under sunlight condition. Journal of Insects as Food and Feed. 10(12):2017–2029. 10.1163/23524588-20230165

Coudron CL, Adamaki-Sotiraki C, Yakti W, et al. 2025. Bugbook: Basic information and good practices on how to maintain stock populations for *Tenebrio molitor* and *Hermetia illucens* for research. 10.1163/23524588-bja10240

Dickerson AJ, Lemke NB, Li C, et al. 2024. Impact of age on the reproductive output of *Hermetia illucens* (Diptera: Stratiomyidae). Crippen T, editor. Journal of Economic Entomology.:toae107. 10.1093/jee/toae107

Generalovic TN, Sandrock C, Roberts BJ, et al. 2023. Cryptic diversity and signatures of domestication in the Black Soldier Fly (Hermetia illucens). :2023.10.21.563413. 10.1101/2023.10.21.563413

Generalovic TN, Zhou W, Zhao LC, et al. 2025. Repeatable phenotypic but not genetic response to selection on body size in the black soldier fly. :2025.02.25.640052. 10.1101/2025.02.25.640052

Giannetti D, Schifani E, Reggiani R, et al. 2022. Do it by yourself: Larval locomotion in the black soldier fly *Hermetia illucens*, with a novel “self-harvesting” method to separate prepupae. Insects. 13(2):127. 10.3390/insects13020127

Harjoko DN, Hua QQH, Toh EMC, et al. 2023. A window into fly sex: mating increases female but reduces male longevity in black soldier flies. Animal Behaviour. 200:25–36. 10.1016/j.anbehav.2023.03.007

Hoffmann L, Hull KL, Bierman A, et al. 2021. Patterns of genetic diversity and mating systems in a mass-reared black soldier fly colony. Insects. 12(6):480. 10.3390/insects12060480

Hoshi T, Higa Y, Chaves LF. 2014. Uranotaenia novobscura ryukyuana (Diptera: Culicidae) Population Dynamics are Denso-Dependent and Autonomous from Weather Fluctuations. Ann Entomol Soc Am. 107(1):136–142. 10.1603/AN13071

James A, Seth A, Marcireau A, et al. 2024. FlyCount: High-speed counting of black soldier flies using neuromorphic sensors. IEEE Sensors Journal.:1–1. 10.1109/JSEN.2024.3504289

Jones BM, Tomberlin JK. 2021. Effects of adult body size on mating success of the black soldier fly, Hermetia illucens (L.) (Diptera: Stratiomyidae). Journal of Insects as Food and Feed. 7(1):5–20. 10.3920/JIFF2020.0001

Kaya C, Generalovic TN, Ståhls G, et al. 2021. Global population genetic structure and demographic trajectories of the black soldier fly, *Hermetia illucens*. BMC Biol. 19(1):94. 10.1186/s12915-021-01029-w

Klüber P, Arous E, Jerschow J, et al. 2024. Fatty acids derived from oviposition systems guide female black soldier flies (*Hermetia illucens*) toward egg deposition sites. Insect Science. 31(4):1231–1248. 10.1111/1744-7917.13287

Klüber P, Arous E, Zorn H, et al. 2023. Protein- and carbohydrate-rich supplements in feeding adult black soldier flies (*Hermetia illucens*) affect life history traits and egg productivity. Life. 13(2):355. 10.3390/life13020355

Kokko H, Klug H, Jennions MG. 2014. Mating systems. In: Shuker DM, Simmons LW, editors. Evoltuion of Insect Mating Systems. Illustrated. United Kingdom: Oxford University Press. p. 78–91.

Laudani F, Campolo O, Latella I, et al. 2024. Does *Hermetia illucens* recognize sibling mates to avoid inbreeding depression? entomologia. 44(5):1225–1232. 10.1127/entomologia/2024/2746

Laursen SF, Flint CA, Bahrndorff S, et al. 2024. Reproductive output and other adult life-history traits of black soldier flies grown on different organic waste and by-products. Waste Management. 181:136–144. 10.1016/j.wasman.2024.04.010

Lemke NB, Dickerson AJ, Tomberlin JK. 2023. No neonates without adults. BioEssays. 45(1):2200162. 10.1002/bies.202200162

Lemke NB, Li C, Dickerson AJ, et al. 2025. Heterogeny in cages: Age-structure and attractant availability impacts fertile egg production in the black soldier fly, Hermetia illucens. Journal of Insects as Food and Feed. Online. 10.1163/23524588-bja10275

Lemke NB, Rollison LN, Tomberlin JK. 2024. Sex-specific perching: Monitoring of artificial plants reveals dynamic female-biased perching behavior in the black soldier fly, *Hermetia illucens* (Diptera: Stratiomyidae). Insects. 15(10):770. 10.3390/insects15100770

Lemke NB, Smith MB, Smink JA, et al. 2025. Wild Flies: Mating Behavior, Adult Foraging, Habitat Use, and Gut Microbiota of Hermetia illucens in Costa Rica. 10.20944/preprints202512.0942.v1

Macklem PT. 2008. Emergent phenomena and the secrets of life. Journal of Applied Physiology. 104(6):1844–1846. 10.1152/japplphysiol.00942.2007

Malawey AS, Mercati D, Love CC, et al. 2019. Adult reproductive tract morphology and spermatogenesis in the black soldier fly (Diptera: Stratiomyidae). Annals of the Entomological Society of America. 112(6):576–586. 10.1093/aesa/saz045

Manas F, Labrousse C, Bressac C. 2025. Plastic responses in sperm expenditure to sperm competition risk in black soldier fly (*Hermetia illucens*, Diptera) males. Journal of Insect Physiology. 161:104751. 10.1016/j.jinsphys.2025.104751

Manas F, Piterois H, Labrousse C, et al. 2024. Gone but not forgotten: dynamics of sperm storage and potential ejaculate digestion in the black soldier fly Hermetia illucens. Royal Society Open Science. 11(10):241205. 10.1098/rsos.241205

Manoukis NC, Collier TC. 2019. Computer vision to enhance behavioral research on insects. Annals of the Entomological Society of America. 112(3):227–235. 10.1093/aesa/say062

Meneguz M, Miranda CD, Cammack JA, et al. 2023. Adult behaviour as the next frontier for optimising industrial production of the black soldier fly *Hermetia illucens* (L.) (Diptera: Stratiomyidae). Journal of Insects as Food and Feed. 9(4):399–414. 10.3920/JIFF2022.0055

Meyermans R, Broeckx L, Mondelaers J, et al. 2025. Exploring the potential of crossbreeding to enhance black soldier fly (Hermetia illucens) production. 10.1163/23524588-00001391

Muraro T, Lalanne L, Pelozuelo L, et al. 2024. Mating and oviposition of a breeding strain of black soldier fly Hermetia illucens (Diptera: Stratiomyidae): polygynandry and multiple egg-laying. Journal of Insects as Food and Feed. 10(8):1–13. 10.1163/23524588-20220175

Nakamura S, Ichiki RT, Shimoda M, et al. 2016. Small-scale rearing of the black soldier fly, *Hermetia illucens* (Diptera: Stratiomyidae), in the laboratory: Low-cost and year-round rearing. Appl Entomol Zool. 51(1):161–166. 10.1007/s13355-015-0376-1

Nawoya S, Ssemakula F, Akol R, et al. 2024. Computer vision and deep learning in insects for food and feed production: A review. Computers and Electronics in Agriculture. 216:108503. 10.1016/j.compag.2023.108503

Normark BB. 2014. Modes of reproduction. In: Shuker DM, Simmons LW, editors. Evoltuion of Insect Mating Systems. Illustrated. United Kingdom: Oxford University Press. p. 78–91.

Oonincx DGAB, Volk N, Diehl JJE, et al. 2016. Photoreceptor spectral sensitivity of the compound eyes of black soldier fly (*Hermetia illucens*) informing the design of LED-based illumination to enhance indoor reproduction. Journal of Insect Physiology. 95:133–139. 10.1016/j.jinsphys.2016.10.006

Park K, Kim W, Lee S, et al. 2010.Seasonal pupation, adult emergence, and mating of black soldier fly, Hermetia illucens (Diptera: Stratiomyidae) in artificial rearing system. :3.

Poh KC, Chaves LF, Reyna-Nava M, et al. 2019. The influence of weather and weather variability on mosquito abundance and infection with West Nile virus in Harris County, Texas, USA. Science of The Total Environment. 675:260–272. 10.1016/j.scitotenv.2019.04.109

Rights JD, Sterba SK. 2020. New recommendations on the use of R-squared differences in multilevel model comparisons. Multivariate Behavioral Research. 55(4):568–599. 10.1080/00273171.2019.1660605

Sharma K, Gold M, Zurbruegg C, et al. 2020. HistoNet: Predicting size histograms of object instances. In: 2020 IEEE Winter Conference on Applications of Computer Vision (WACV). Snowmass Village, CO, USA: IEEE. p. 3626–3634. 10.1109/WACV45572.2020.9093484

Shuker DM, Simmons LW, editors. 2014. The evolution of insect mating systems. University of Oxford: Oxford University Press.

Shumway RH, Stoffer DS. 2017a. Time series analysis and Its applications: With R examples. Cham: Springer International Publishing (Springer Texts in Statistics). 10.1007/978-3-319-52452-8

Shumway RH, Stoffer DS. 2017b. ARIMA models. In: Shumway RH, Stoffer DS, editors. Time Series Analysis and Its Applications: With R Examples. Cham: Springer International Publishing. p. 75–163. 10.1007/978-3-319-52452-8_3

Smetana S, Coudron C, Deruytter D, et al. 2025. BugBook: Data analysis methods in studies of insects for food and feed. 10.1163/23524588-bja10209

[Software] Wei T, Simko V. 2024. R package “corrplot”: Visualization of a correlation matrix. https://github.com/taiyun/corrplot.

Thinn AA, Kainoh Y. 2022. Effect of diet on the longevity and oviposition performance of black soldier flies, Hermetia illucens (Diptera: Stratiomyidae). Japan Agricultural Research Quarterly: JARQ. 56(2):211–217. 10.6090/jarq.56.211

Tomberlin JK, Klammsteiner T, Lemke N, et al. 2025. BugBook: Black soldier fly as a model to assess behaviour of insects mass produced as food and feed. 10.1163/23524588-bja10225

Tomberlin JK, Sheppard DC. 2001. Lekking behavior of the black soldier fly (Diptera: Stratiomyidae). Florida Entomologist. 84(4):729–730.

Tomberlin JK, Sheppard DC. 2002. Factors influencing mating and oviposition of black soldier flies (Diptera: Straiomyidae) in a colony. Journal of Entomological Science. 37(4):345–352. 10.18474/0749-8004-37.4.345

Tomberlin JK, Sheppard DC, Joyce JA. 2002. Selected life-history traits of black soldier flies (Diptera: Stratiomyidae) reared on three artificial diets. an. 95(3):379–386. 10.1603/0013-8746(2002)095%5B0379:SLHTOB%5D2.0.CO;2

Zhang J, Huang L, He J, et al. 2010. An artificial light source influences mating and oviposition of black soldier flies, Hermetia illucens. Journal of Insect Science. 10(202):1–7. 10.1673/031.010.20201

