## Supplementary Info for "Temporal Trends: Phase-shifted time-series analysis reveals highly correlated reproductive behaviors in the black soldier fly, *Hermetia illucens* (Diptera: Stratiomyidae)"

`Noah B Lemke, PhD
4921 Crawford St 3

Houston, Texas, USA 77004

Phone: US +1 630 347 3302

RESEARCH ARTICLE: Temporal Trends: Phase-shifted time-series analysis reveals highly correlated reproductive behaviors in the black soldier fly, *Hermetia illucens* (Diptera: Stratiomyidae)

AUTHORS

Noah B Lemke 1 2, Chujun Li 3, Jeroen De Smet 2

AFFILIATIONS

1 - Department of Entomology, 370 Olsen Blvd Suite 412, Texas A&M University, College Station, Texas, USA 77843

2 - Research Group for Insect Production and Processing, Department of Microbial and Molecular Systems, KU Leuven, Campus Geel, Kleinhoefstraat 4, 2440 Geel, Belgium

3 - Guangzhou Unique Biotechnology, Room 1717, 101 Science Avenue, Huanpu District, Guangzhou, Guangdong, China

ENGLISH ABSTRACT (213 words)

For those researching reproduction of the black soldier fly (BSF), finding meaningful correlations between mating and downstream reproductive fitness metrics, owing to complex reproductive behavior and physiology, has been challenging. On the one hand, mating is not necessarily deterministic of fertile eggs being produced, since females might lay unfertile eggs both before and after mating. But conversely, the process is not ‘random’ either, since fertility is predicated on the exchange of gametes during mating. To investigate the relationship between mating and other reproductive metrics, a cross-correlation analysis between lagged time series was conducted on data previously collected during a semi-outdoor greenhouse experiment. This process compared the average percent correlation between un-lagged mating with oviposition in traps, oviposition outside traps, total weight of eggs deposited, and hatch rate, which were lagged 0 to 6 days. This revealed that average mating was 97.54% correlated with oviposition in traps lagged by 2 days. For oviposition outside of traps, total egg weights, and hatch rate, these were also ~71.63% correlated, 60.29% correlated, and 75.16% correlated with mating respectively, which were optimized when the time series were lagged n = 3 days. These trends likely have a biological basis since flies decrease their reproductive activity as they expend energy and senesce, aligning with temporal patterns consistently observed during BSF rearing.

Generalovic TN, Sandrock C, Roberts BJ, et al. 2023. Cryptic diversity and signatures of domestication in the Black Soldier Fly (Hermetia illucens). :2023.10.21.563413. https://doi.org/10.1101/2023.10.21.563413

Generalovic TN, Zhou W, Zhao LC, et al. 2025. Repeatable phenotypic but not genetic response to selection on body size in the black soldier fly. :2025.02.25.640052. https://doi.org/10.1101/2025.02.25.640052

Giannetti D, Schifani E, Reggiani R, et al. 2022. Do it by yourself: Larval locomotion in the black soldier fly *Hermetia illucens*, with a novel “self-harvesting” method to separate prepupae. Insects. 13(2):127. https://doi.org/10.3390/insects13020127

Harjoko DN, Hua QQH, Toh EMC, et al. 2023. A window into fly sex: mating increases female but reduces male longevity in black soldier flies. Animal Behaviour. 200:25–36. https://doi.org/10.1016/j.anbehav.2023.03.007

Hoffmann L, Hull KL, Bierman A, et al. 2021. Patterns of genetic diversity and mating systems in a mass-reared black soldier fly colony. Insects. 12(6):480. https://doi.org/10.3390/insects12060480

Hoshi T, Higa Y, Chaves LF. 2014. Uranotaenia novobscura ryukyuana (Diptera: Culicidae) Population Dynamics are Denso-Dependent and Autonomous from Weather Fluctuations. Ann Entomol Soc Am. 107(1):136–142. https://doi.org/10.1603/AN13071

James A, Seth A, Marcireau A, et al. 2024. FlyCount: High-speed counting of black soldier flies using neuromorphic sensors. IEEE Sensors Journal.:1–1. https://doi.org/10.1109/JSEN.2024.3504289

Jones BM, Tomberlin JK. 2021. Effects of adult body size on mating success of the black soldier fly, *Hermetia illucens* (L.) (Diptera: Stratiomyidae). Journal of Insects as Food and Feed. 7(1):5–20. https://doi.org/10.3920/JIFF2020.0001

Kaya C, Generalovic TN, Ståhls G, et al. 2021. Global population genetic structure and demographic trajectories of the black soldier fly, *Hermetia illucens*. BMC Biol. 19(1):94. https://doi.org/10.1186/s12915-021-01029-w

Klüber P, Arous E, Jerschow J, et al. 2024. Fatty acids derived from oviposition systems guide female black soldier flies (*Hermetia illucens*) toward egg deposition sites. Insect Science. 31(4):1231–1248. https://doi.org/10.1111/1744-7917.13287

Klüber P, Arous E, Zorn H, et al. 2023. Protein- and carbohydrate-rich supplements in feeding adult black soldier flies (*Hermetia illucens*) affect life history traits and egg productivity. Life. 13(2):355. https://doi.org/10.3390/life13020355

Kokko H, Klug H, Jennions MG. 2014. Mating systems. In: Shuker DM, Simmons LW, editors. Evoltuion of Insect Mating Systems. Illustrated. United Kingdom: Oxford University Press. p. 78–91.

Laudani F, Campolo O, Latella I, et al. 2024. Does *Hermetia illucens* recognize sibling mates to avoid inbreeding depression? entomologia. 44(5):1225–1232. https://doi.org/10.1127/entomologia/2024/2746

Laursen SF, Flint CA, Bahrndorff S, et al. 2024. Reproductive output and other adult life-history traits of black soldier flies grown on different organic waste and by-products. Waste Management. 181:136–144. https://doi.org/10.1016/j.wasman.2024.04.010

Lemke NB, Dickerson AJ, Tomberlin JK. 2023. No neonates without adults. BioEssays. 45(1):2200162. https://doi.org/10.1002/bies.202200162

Lemke NB, Li C, Dickerson AJ, et al. 2025. Heterogeny in cages: Age-structure and attractant availability impacts fertile egg production in the black soldier fly, *Hermetia illucens*. Journal of Insects as Food and Feed. Online. https://doi.org/10.1163/23524588-bja10275

Lemke NB, Rollison LN, Tomberlin JK. 2024. Sex-specific perching: Monitoring of artificial plants reveals dynamic female-biased perching behavior in the black soldier fly, *Hermetia illucens* (Diptera: Stratiomyidae). Insects. 15(10):770. https://doi.org/10.3390/insects15100770

Lemke NB, Smith MB, Smink JA, et al. 2025. Wild Flies: Mating Behavior, Adult Foraging, Habitat Use, and Gut Microbiota of *Hermetia illucens* in Costa Rica. https://doi.org/10.20944/preprints202512.0942.v1

Macklem PT. 2008. Emergent phenomena and the secrets of life. Journal of Applied Physiology. 104(6):1844–1846. https://doi.org/10.1152/japplphysiol.00942.2007

Malawey AS, Mercati D, Love CC, et al. 2019. Adult reproductive tract morphology and spermatogenesis in the black soldier fly (Diptera: Stratiomyidae). Annals of the Entomological Society of America. 112(6):576–586. https://doi.org/10.1093/aesa/saz045

Manas F, Labrousse C, Bressac C. 2025. Plastic responses in sperm expenditure to sperm competition risk in black soldier fly (*Hermetia illucens*, Diptera) males. Journal of Insect Physiology. 161:104751. https://doi.org/10.1016/j.jinsphys.2025.104751

Manas F, Piterois H, Labrousse C, et al. 2024. Gone but not forgotten: dynamics of sperm storage and potential ejaculate digestion in the black soldier fly Hermetia illucens. Royal Society Open Science. 11(10):241205. https://doi.org/10.1098/rsos.241205

Manoukis NC, Collier TC. 2019. Computer vision to enhance behavioral research on insects. Annals of the Entomological Society of America. 112(3):227–235. https://doi.org/10.1093/aesa/say062

Meneguz M, Miranda CD, Cammack JA, et al. 2023. Adult behaviour as the next frontier for optimising industrial production of the black soldier fly *Hermetia illucens* (L.) (Diptera: Stratiomyidae). Journal of Insects as Food and Feed. 9(4):399–414. https://doi.org/10.3920/JIFF2022.0055

Meyermans R, Broeckx L, Mondelaers J, et al. 2025. Exploring the potential of crossbreeding to enhance black soldier fly (Hermetia illucens) production. https://doi.org/10.1163/23524588-00001391

Muraro T, Lalanne L, Pelozuelo L, et al. 2024. Mating and oviposition of a breeding strain of black soldier fly Hermetia illucens (Diptera: Stratiomyidae): polygynandry and multiple egg-laying. Journal of Insects as Food and Feed. 10(8):1–13. https://doi.org/10.1163/23524588-20220175

Nakamura S, Ichiki RT, Shimoda M, et al. 2016. Small-scale rearing of the black soldier fly, *Hermetia illucens* (Diptera: Stratiomyidae), in the laboratory: Low-cost and year-round rearing. Appl Entomol Zool. 51(1):161–166. https://doi.org/10.1007/s13355-015-0376-1

Nawoya S, Ssemakula F, Akol R, et al. 2024. Computer vision and deep learning in insects for food and feed production: A review. Computers and Electronics in Agriculture. 216:108503. https://doi.org/10.1016/j.compag.2023.108503

Normark BB. 2014. Modes of reproduction. In: Shuker DM, Simmons LW, editors. Evoltuion of Insect Mating Systems. Illustrated. United Kingdom: Oxford University Press. p. 78–91.

Oonincx DGAB, Volk N, Diehl JJE, et al. 2016. Photoreceptor spectral sensitivity of the compound eyes of black soldier fly (*Hermetia illucens*) informing the design of LED-based illumination to enhance indoor reproduction. Journal of Insect Physiology. 95:133–139. https://doi.org/10.1016/j.jinsphys.2016.10.006

Park K, Kim W, Lee S, et al. 2010. Seasonal pupation, adult emergence, and mating of black soldier fly, *Hermetia illucens* (Diptera: Stratiomyidae) in artificial rearing system. :3.

Poh KC, Chaves LF, Reyna-Nava M, et al. 2019. The influence of weather and weather variability on mosquito abundance and infection with West Nile virus in Harris County, Texas, USA. Science of The Total Environment. 675:260–272. https://doi.org/10.1016/j.scitotenv.2019.04.109

Rights JD, Sterba SK. 2020. New recommendations on the use of R-squared differences in multilevel model comparisons. Multivariate Behavioral Research. 55(4):568–599. https://doi.org/10.1080/00273171.2019.1660605

Sharma K, Gold M, Zurbruegg C, et al. 2020. HistoNet: Predicting size histograms of object instances. In: 2020 IEEE Winter Conference on Applications of Computer Vision (WACV). Snowmass Village, CO, USA: IEEE. p. 3626–3634. https://doi.org/10.1109/WACV45572.2020.9093484

Shuker DM, Simmons LW, editors. 2014. The evolution of insect mating systems. University of Oxford: Oxford University Press.

Shumway RH, Stoffer DS. 2017a. Time series analysis and Its applications: With R examples. Cham: Springer International Publishing (Springer Texts in Statistics). https://doi.org/10.1007/978-3-319-52452-8

Shumway RH, Stoffer DS. 2017b. ARIMA models. In: Shumway RH, Stoffer DS, editors. Time Series Analysis and Its Applications: With R Examples. Cham: Springer International Publishing. p. 75–163. https://doi.org/10.1007/978-3-319-52452-8_3

Smetana S, Coudron C, Deruytter D, et al. 2025. BugBook: Data analysis methods in studies of insects for food and feed. https://doi.org/10.1163/23524588-bja10209

[Software] Wei T, Simko V. 2024. R package “corrplot”: Visualization of a correlation matrix. https://github.com/taiyun/corrplot.

Thinn AA, Kainoh Y. 2022. Effect of diet on the longevity and oviposition performance of black soldier flies, Hermetia illucens (Diptera: Stratiomyidae). Japan Agricultural Research Quarterly: JARQ. 56(2):211–217. https://doi.org/10.6090/jarq.56.211

Tomberlin JK, Klammsteiner T, Lemke N, et al. 2025. BugBook: Black soldier fly as a model to assess behaviour of insects mass produced as food and feed. https://doi.org/10.1163/23524588-bja10225

Tomberlin JK, Sheppard DC. 2001. Lekking behavior of the black soldier fly (Diptera: Stratiomyidae). Florida Entomologist. 84(4):729–730.

Tomberlin JK, Sheppard DC. 2002. Factors influencing mating and oviposition of black soldier flies (Diptera: Straiomyidae) in a colony. Journal of Entomological Science. 37(4):345–352. https://doi.org/10.18474/0749-8004-37.4.345

Tomberlin JK, Sheppard DC, Joyce JA. 2002. Selected life-history traits of black soldier flies (Diptera: Stratiomyidae) reared on three artificial diets. an. 95(3):379–386. https://doi.org/10.1603/0013-8746(2002)095%5B0379:SLHTOB%5D2.0.CO;2
